# Correspondence: Recurrent *GNAQ* mutation encoding T96S in natural killer/T cell lymphoma

**DOI:** 10.1101/2020.04.01.020198

**Authors:** Jing Quan Lim, Soon Thye Lim, Choon Kiat Ong

**Author notes:** Corresponding authors: Jing Quan Lim and Choon Kiat Ong.

## Abstract

Recurrent *GNAQ* mutation encoding p.T96S in natural-killer/T cell lymphoma (NKTCL) was recently reported in 8.7% (11/127) of NKTCL patients. At the time of publication, this phenomenon was not observed in preceding genomic studies of NKTCL. We suspected that p.T96S was a false-positive somatic call due to misaligned sequencing reads that originated from the highly similar GNAQ-pseudogene (*GNAQP*) chr2q21.1 locus. Linkage disequilibrium analysis also revealed that *GNAQP* has high frequencies of co-occurring polymorphic mismatches which led to the preferential misalignment of sequencing reads to the *GNAQ* chr9q21.2 locus instead. This correspondence implicates our interpretation of true-positive somatic variants and many other studies which could be affected by similar suboptimal interpretation of somatic mutations.

## Introduction

Next-generation sequencing (NGS) has enabled the interrogation of DNA sequences at an unprecedented fashion. After the sequencing of genomic library DNA, all reference-based bioinformatics analysis would involve an mandatory ‘alignment’ step before many downstream analysis could take place. A bioinformatics tool, such as BWA^1^, would handle this ‘alignment’ step and report the positional coordinates of each NGS read with respect to the reference genome that it has based the alignments on (Fig. 1a). An aligner would score each seed alignment, by accounting for the matches, mismatches or gaps with a scoring function (Fig. 1b), between the read and the locality of the reference genome that the aligner assigns it to.

**Fig. 1.**
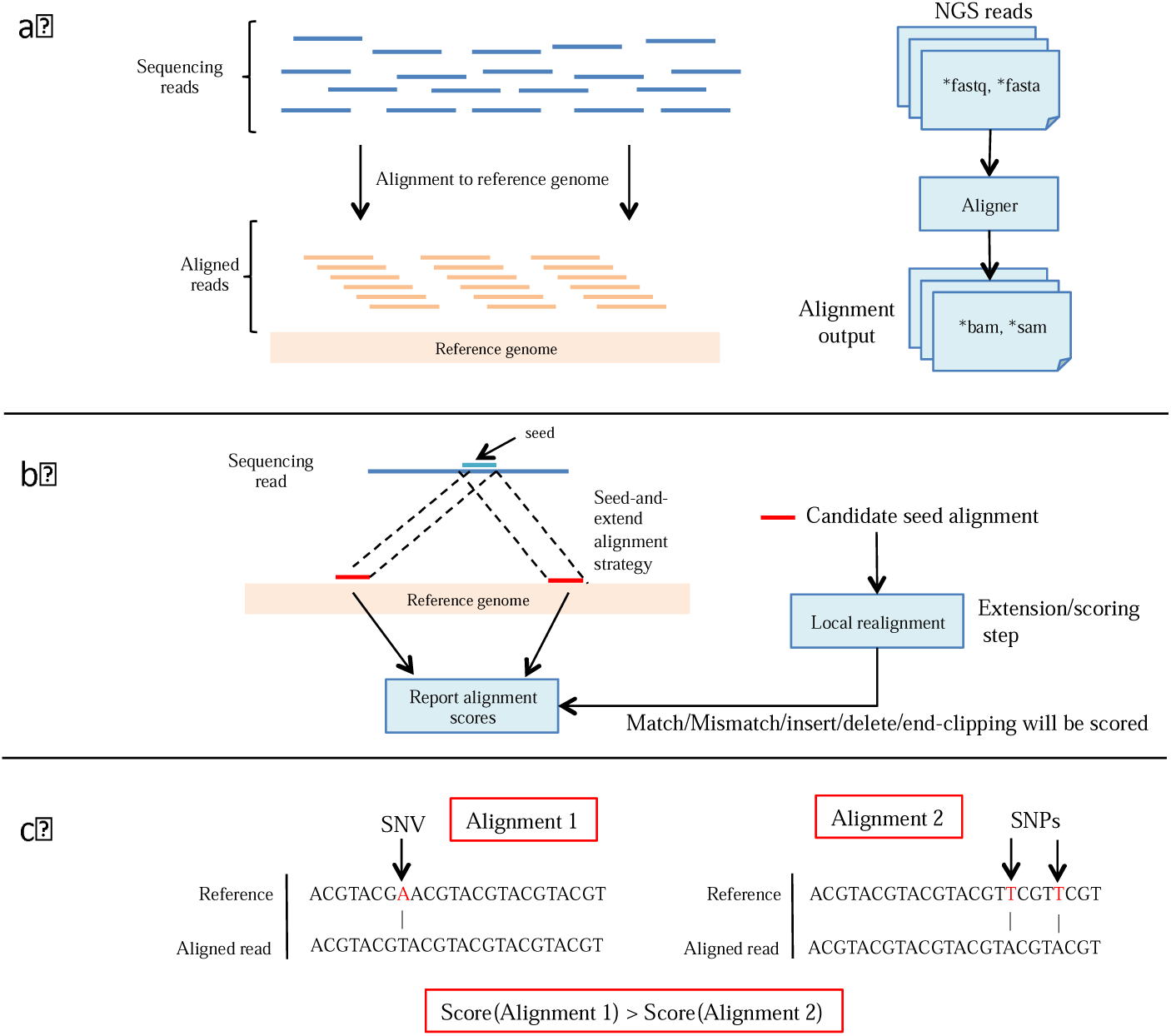
Alignment of sequencing reads to a reference genome. Panel A shows the bioinformatics step of alignment where an aligner finds the positional origin of sequencing reads in a reference genome. Panel B shows the seed-and-extend strategy employed by many aligners to align reads onto a reference genome. Panel C shows a possible senario whereby a 1-mismatch alignment (left) is returned as the best hit instead of a correct 2-SNPs alignment (right).

In practice, the seed-extended alignment with the highest score would be the primary alignment for a read. However, the primary alignment might not always be correct for a read. For instance, a single-nucleotide polymorphism (SNP) would cause a mismatch in the alignment between a read and the reference genome and will not be considered as an exact-matching alignment instead (Fig. 1c). As such, a correct alignment covering common polymorphisms would not be considered as a ‘better’ hit, if another incorrect alignment containing fewer mismatches would to be found by the aligner.

Formalin-fixed paraffin-embedded (FFPE) archival materials present great opportunities to study various diseases. However, FFPE DNA are often more fragmented and will yield shorter NGS reads as compared to their Fresh/Frozen (FF) counterparts. In general, a shorter read-length would contain less information content for a read to be aligned uniquely and would be misaligned more often than NGS reads of longer read-lengths. As such, subsequent analysis of misaligned SNP-stricken NGS-reads would cascade into a mirage of results.

As Zhaoming Li *et al*. has used FFPE materials for all their sequencing work, we hope to find out of if the recurrent *GNAQ* mutations encoding p.T96S and p.Y101X are real. Below, we present our case and possible solutions to further confirm if the said somatic mutations are real.

### Summary of Zhaoming Li *et al*., *Nature Communications* 2019

A recent study found recurrent *GNAQ* mutation encoding p.T96S in 8.7% (11/127) of natural-killer/T cell lymphoma (NKTCL) using NGS technologies^2^. The paper demonstrated that GNAQ deficiency led to enhanced NK cell survival in conditional knockout mice (*Ncr1-Cre-Gnaq*^*fl/fl*^) via the inhibition of AKT and MAPK signaling pathways. It was also shown to be clinically important as patients with GNAQ p.T96S have inferior survival and could be relevant for the development of therapies.

### Recurrent p.T96S and p.Y101X Somatic Mutations are Potential Artefacts from Bioinformatics Misalignments

It was of peculiar interest to us that the two *GNAQ* hotspot somatic mutations (p.T96S and p.Y101X) reported in this study were never reported in other NKTCL studies that used NGS too^3-6^. We analyzed the Sanger sequences provided in Supplementary Fig. 4 of the work in question and realized that the single-nucleotide variant (SNV) that encoded for p.T96S had a minor allele frequency (MAF) of 1.18% (1386/117782, ExAC^7^ and gnomAD^8^ databases; dbSNP151, rs753716491), which we found to be too common if it was to contribute sustantially to the pathogenesis of NKTCL. Moreover, the authors wrote in the published work that the *GNAQ* somatic mutations encoding for p.Y101X tends to co-occur with p.T96S. However, the *GNAQ* somatic mutation that encoded for p.Y101X was not marked as a common SNP by germline databases and it was also functionally redundant for a stop-gain (p.Y101X) mutation to co-occur with another missense (p.T96S) mutation on the same gene. This was the first hint that the alignments to the *GNAQ* locus that encoded for both p.T96S and p.Y101X were erroneous.

Next, we wanted to further analyze the NGS reads that encoded for both p.T96S and p.Y101X somatic mutations were indeed misaligned. We simulated 100 bp long NGS reads that would encode for both p.T96S and p.Y101X somatic mutations from the genomic locus of *GNAQ* using the same hg19 reference that the authors have used and realigned the *in silico* reads back to the same reference (Fig. 2a). The reads were multi-mapped to the genomic loci of *GNAQ* and *GNAQ*-psuedogene-1 (*GNAQP*) at chr9q21.2 and chr2q21.1, respectively. As expected, the read was realigned back to the *GNAQ* locus that it was simulated from and recapitulated the two simulated SNVs too; chr9:80537095[G>T] (p.Y101X) and chr9:80537112[T>A] (p.T96S, rs753716491). Next, Fig. 2b shows that the realignment mapped the read to *GNAQP* too and yielded three SNVs, all of which are common SNPs as denoted by their respective dbSNP IDs; chr2:132182138[G>T] (rs3730150), chr2:132182159[T>C] (rs3730148) and chr2:132182199[C>T] (rs3730153).

**Fig. 2.**
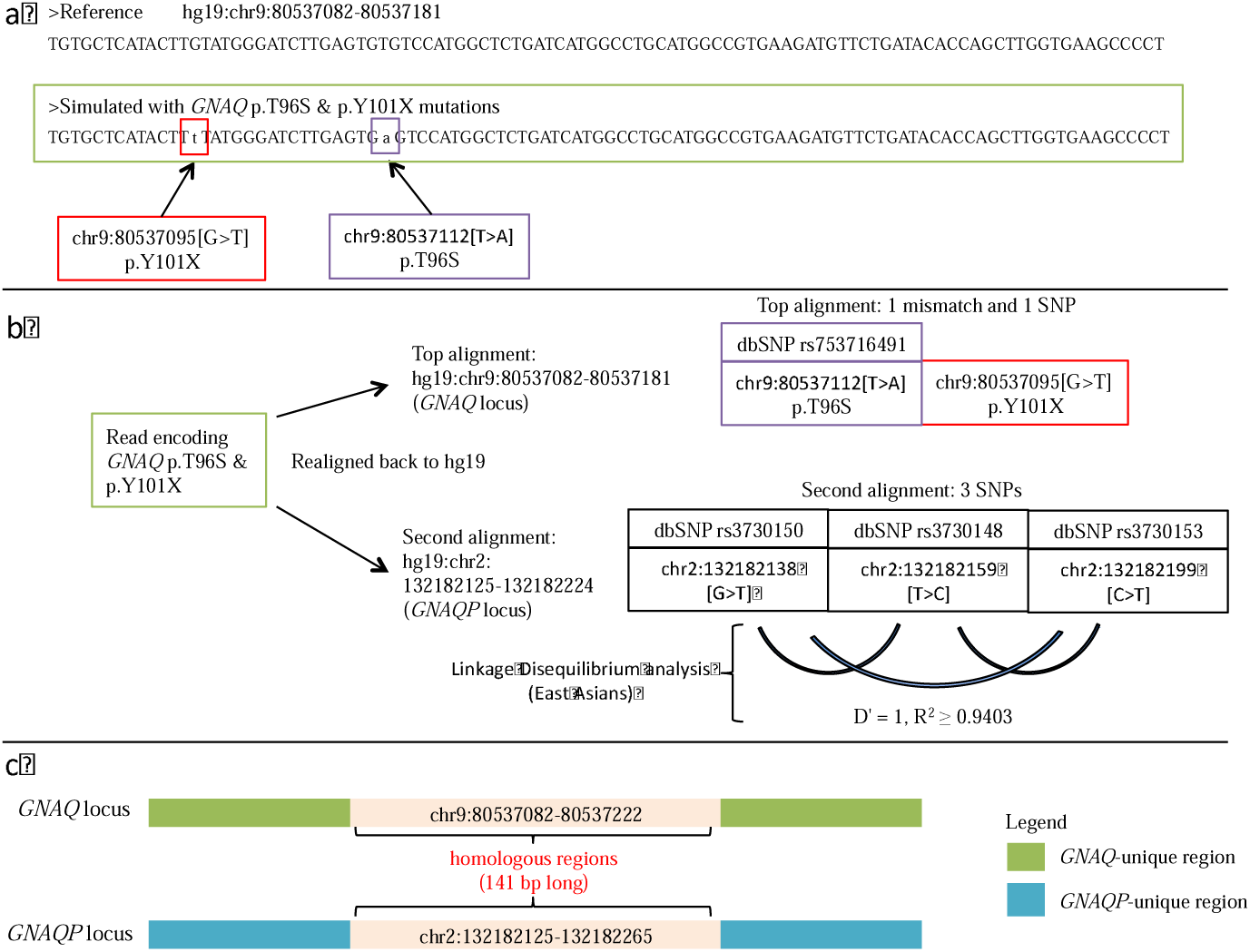
GNAQ p.T96S and p.Y101X mutations could be the results of misaligned sequencing reads from GNAQ-Pseudogene-1. Panel A shows an *in-silico* simulated read that would encode for GNAQ p.T96S and p.101X mutations. Panel B shows the top alignments of the read that would encode for GNAQ p.T96S and p.101X. The read aligns to both *GNAQ* (with 1 mismatch and 1 SNP) and *GNAQP* (with 3 SNPs) simultaneously. Linkage disequilibirum analysis of the 3 SNPs from the *GNAQP* locus also showed that they tend to co-occur and cause an misalignment to *GNAQ* locus. This misalignment would yield the wrong callings of p.T96S and p.101X mutations. Panel C describes the *GNAQ*-*GNAQP* homologous regions that implicated p.T96S and p.Y101X, and rs3730150, rs3730148 and rs3730153 in the *GNAQ* and *GNAQP* loci, respectively. The immediate regions outside of chr9:80537082-80537173 are unique to *GNAQ* that would further help Zhaoming Li, *et al*. to further validate their current findings.

We performed linkage disequilibrium (LD - LDlink) analysis^9^ of all three possible pairwise combinations of the three SNPs within *GNAQP* and found that they were extremely likely to co-occur together as a triplet of SNPs within *GNAQP* (Fig. 2b, D’ = 1, R^2^ ≥ 0.9403). As such, NGS reads that were representing these SNPs would be misaligned to *GNAQ* instead and be misinterpreted for somatic mutations encoding for p.T96S and p.Y101X instead.

### Additional Checks to Confirm the Validity of *GNAQ* somatic mutations encoding for p.T96S and p.Y101X

By performing an modified SNP-aware version of Smith-Waterman alignment^10^ between the genomic sequences of *GNAQ* and *GNAQP*, we found that chr9:80537082-80537222 and chr2:132182125-132182265 were homologous and encapulated all the SNPs and variants that implicated the validity of the reported *GNAQ* somatic mutations (Fig. 2c). To confirm the reported mutations, the authors need to look for alignments that can satisfy the following two criteria simultaneously. 1) The alignment must represent *GNAQ* mutations that encode for p.T96S and p.Y101X. 2) The alignment must extend errorlessly beyond chr9:80537082-80537222. If either of the two criteria cannot be satisfied, then the validity of the reported *GNAQ* somatic mutations in NKTCL cannot stand.

With the general implication that our combined findings would bring about, we might have to take a step back and revisit *GNAQ* p.T96S, p.Y101X and rs753716491 non-reference alleles that have also been reported, but unmentioned studies, in this correspondence.

## Conclusion

For the 127 NKTCLs that were studied by Zhaoming Li *et al*., they were all FFPE archival materials and 101 of them had matched whole blood as its germline counterpart. DNA extracted from whole-blood are typically less fragmented and tends to yield longer NGS read-lengths than DNA extracted from FFPE archival materials. This allows NGS reads sequenced from whole-blood to align more accurately than the ones sequenced from FFPE archival materials onto a reference genome. This would mean that sequencing reads that originated from one genomic locus could be mapped to more than one genomic loci and yielded variant artefacts in subsequent downstream analyses.

In an analysis for somatic mutations, the germline mutations would be subtracted from the tumor mutations. In this case, the *GNAQ* p.T96S and p.Y101X somatic artefacts leaked through the subtraction step as reads sequenced from the GNAQ and GNAQP loci were aligned differently from both FFPE archival tumor and normal whole-nlood samples. It was an ill-fated combination of the following three criteria 1) Short tumor reads that failed to align correctly 2) Long germline reads that aligned correctly and 3) SNP-stricken genomic region from where the tumor reads were sequenced that yielded the *GNAQ* p.T96S and p.Y101X artefacts. As such, we strongly believe that it was purely unintentional that led to this case.

## References

1 Li, H. & Durbin, R. Fast and accurate short read alignment with Burrows-Wheeler transform. Bioinformatics 25, 1754–1760, doi:10.1093/bioinformatics/btp324 (2009).

2 Li, Z. et al. Recurrent GNAQ mutation encoding T96S in natural killer/T cell lymphoma. Nature communications 10, doi:10.1038/s41467-019-12032-9 (2019).

3 Koo, G. C. et al. Janus kinase 3-activating mutations identified in natural killer/T-cell lymphoma. Cancer discovery 2, 591–597, doi:10.1158/2159-8290.CD-12-0028 (2012).

4 Jiang, L. et al. Exome sequencing identifies somatic mutations of DDX3X in natural killer/T-cell lymphoma. Nature genetics 47, 1061–1066, doi:10.1038/ng.3358 (2015).

5 Song, T. L. et al. Oncogenic activation of STAT3 pathway drives PD-L1 expression in natural killer/T cell lymphoma. Blood, blood-2018-2001-829424, doi:10.1182/blood-2018-01-829424 (2018).

6 Wen, H. et al. Recurrent ECSIT mutation encoding V140A triggers hyperinflammation and promotes hemophagocytic syndrome in extranodal NK/T cell lymphoma. Nature Medicine 24, 154–164, doi:10.1038/nm.4456 (2018).

7 Lek, M. et al. Analysis of protein-coding genetic variation in 60,706 humans. Nature 536, 285–291, doi:10.1038/nature19057 (2016).

8 Karczewski, K. J. et al., doi:10.1101/531210 (2019).

9 Machiela, M. J. & Chanock, S. J. LDlink: a web-based application for exploring population-specific haplotype structure and linking correlated alleles of possible functional variants. Bioinformatics 31, 3555–3557, doi:10.1093/bioinformatics/btv402 (2015).

10 Smith, T. F. & Waterman, M. S. Identification of common molecular subsequences. Journal of Molecular Biology 147, 195–197, doi:10.1016/0022-2836(81)90087-5 (1981).

